# Metabolic Reprogramming Coordinates Mannose and Glutamine Metabolism to Maintain Glucose Homeostasis During Glycosuria

**DOI:** 10.64898/2026.05.20.726580

**Authors:** Nadia Rashid, Moses Otunla, Nazmul Hasan, Michael J. Hodges, Heba H. Qaissi, Tumininu S. Faniyan, Pontiana Ritika Clement, Penghui Lin, Mohamed M. Y. Kaddah, Teresa A. Cassel, Donald A. Morgan, Kamal Rahmouni, Kavaljit H. Chhabra

## Abstract

Glycosuria, whether genetically induced or triggered by SGLT2 inhibitors, activates compensatory glucose-producing pathways that limit glucose lowering in type 2 diabetes. To define these pathways, we studied renal *Glut2* knockout mice, which progressively lose *Slc5a2* (encoding SGLT2) expression yet maintain normoglycemia despite marked urinary glucose loss. Metabolic profiling and isotope tracing revealed coordinated adaptations in mannose and glutamine metabolism during glycosuria. Skeletal muscle reduced glucose utilization and instead oxidized mannose, while whole-body glycolysis declined, establishing a systemic glucose-sparing state. Disruption of glutamine transport or mannose utilization caused hypoglycemia in mice treated with an SGLT2 inhibitor, demonstrating dependence on these substrates to maintain glucose homeostasis during glycosuria. Multiomic profiling revealed increased expression and chromatin accessibility of mannose and glutamine transport pathways. These findings identify a kidney-driven metabolic program that preserves systemic glucose homeostasis during glycosuria and may inform strategies to optimize the glucose-lowering efficacy of SGLT2 inhibitors.

## Introduction

The kidney regulates systemic glucose homeostasis by reabsorbing filtered glucose through transporters such as GLUT2 and SGLT2. When this process is disrupted, large amounts of glucose are lost in the urine (renal glycosuria), yet blood glucose levels often remain remarkably stable - a paradox that highlights unknown compensatory mechanisms. Genetic deficiency of *GLUT2*^*1,2*^ or *SGLT2*^*3*^ causes renal glycosuria, resulting in high urinary glucose excretion despite normal blood glucose. Similarly, SGLT2 inhibitors increase glucose excretion in urine and are used to manage type 2 diabetes^4^. However, their glucose lowering efficacy is attenuated^5^ in part due to endogenous glucose compensation that offsets the glucose loss in urine.

Consistent with this concept, healthy individuals^6^ and rodents^7^ treated with SGLT2 inhibitors maintain normal blood glucose levels despite massive glycosuria. A similar phenotype is observed in mice lacking renal *Glut2*^2^ or *Sglt2*^8^, which maintain normoglycemia through compensatory glucose production. Previous studies suggest that central mechanisms contribute to this response: the hypothalamus regulates glucose reabsorption^9-11^ through renal GLUT2 and promotes glucose production^12^ during glycosuria. Renal nerves have also been implicated in this compensatory response, although their contribution appears to be partial and context dependent^12-14^. For example, renal denervation reduces compensatory glucose production by only ∼50% in renal *Glut2* knockout (r*Glut2* KO) mice^12^, indicating that additional mechanisms are responsible for glucose compensation. Similarly, renal denervation in human subjects treated with SGLT2 inhibitors does not affect compensatory glucose production^13^. By contrast, glucose production in response to SGLT2 inhibitors is attenuated in another study involving kidney transplantation and bilateral nephrectomy^14^. Thus, the metabolic pathways that sustain glucose homeostasis during chronic renal glycosuria remain undefined, and identifying these adaptations may reveal strategies to enhance the glucose-lowering efficacy of SGLT2 inhibitors in type 2 diabetes.

To investigate systemic adaptations to chronic renal glucose loss, we studied mouse models of renal glycosuria. These include r*Glut2* KO mice ^2^, which develop progressive loss of *Slc5a2* expression (encoding SGLT2) and marked urinary glucose excretion while maintaining normal blood glucose levels. This phenotype has been independently confirmed and reviewed by other research groups^15,16^, validating r*Glut2* KO mice as a genetic model of sustained renal glycosuria. We also examined mice treated with the SGLT2 inhibitor dapagliflozin to assess whether similar metabolic adaptations occur during pharmacologically induced glycosuria.

Here, we identify coordinated regulation of mannose and glutamine metabolism as key components of a kidney -driven metabolic program that maintains systemic glucose homeostasis during persistent glycosuria.

## Methods

### Untargeted primary metabolism and stable isotope-resolved metabolomics (SIRM) analysis

r*Glut2* KO mice and their control littermates^2,12^ were used in this study. We have described the production, genotyping of r*Glut2* KO and their control mice previously^2^. After genotyping, mice were randomly assigned to different experimental groups including random order of treatments and measurements. The randomization was performed using GraphPad Prism 10.2.3. Researchers were not blinded to group allocation during the experiment because of obvious phenotypic differences (such as polyuria and polydipsia) between the control and experimental groups of mice. The age and sex of mice included in each experiment in this study are mentioned in the figure legends. No mice or data points were excluded from the analysis. We did not repeat all the experiments in both sexes because our previous study^2,12^ demonstrated that male and female r*Glut2* KO mice exhibited a similar phenotype compared to their corresponding control groups in the context of glucose homeostasis. All mouse studies were approved by University of Kentucky, University of Iowa, or University of Massachusetts.

Primary metabolism in mouse kidneys was analyzed using GC-TOF-MS (West coast metabolomics center, UC Davis). The kidney samples were collected around 9am from non-fasted male mice and flash frozen in liquid nitrogen. The samples were then processed for metabolism analysis as described previously^17^. For SIRM analysis, ^13^C_6_ glucose (CLM-1396-0.25, Cambridge Isotope Laboratories, USA) or ^13^C_6_ mannose (CLM-6567-0.25, Cambridge Isotope Laboratories, USA) was administered by oral gavage (60 mg/mouse, the isotopes were dissolved in water with the concentration of 200 mg/ml and 300 µl was administered to each mouse) to separate cohorts of mice. The rationale for giving equal amount of glucose or mannose to each mouse has been described previously^9^. The mice were euthanized by decapitation 60 min. after the stable isotope administration to collect trunk blood and tissues in this order: urine, trunk blood, kidney, liver, and skeletal muscle (all samples were collected within six min. of euthanizing the mice). The samples were flash frozen in liquid nitrogen, stored at -80^0^C until analyzed as described previously^18^. Ion chromatography coupled with ultra-high resolution Fourier transform mass spectrometry (IC-UHR-FTMS) was used to quantify glucose and mannose metabolites. The frozen tissue samples were pulverized to a fine powder under liquid nitrogen prior to 3-phase extraction. Polar/non-polar metabolites and proteins were extracted simultaneously using acetonitrile:H2O:chloroform (2:1.5:1, v/v) solvent partitioning method as described previously^19^.

IC-UHR-FTMS was carried out as described previously^20^. Metabolites were separated on a Thermo Scientific Dionex™ IonPac AG11-HC analytical column paired with a Dionex IonPac AS11-HC guard column in a Dionex ICS-5000+ DP Ion Chromatography system interfaced to a Thermo Fisher Orbitrap Fusion™ Tribrid™ mass spectrometer (Waltham, MA, USA) run in negative MS1 mode at a resolving power of 500,000 at m/z 200 and an m/z range of 80 to 800. Data were analyzed using TraceFinder™ software and an in-house database. MS1 peak areas of assigned metabolites and isotopologues were corrected for natural abundance and quantified against a calibration standard mixture. Metabolite amounts thus obtained were normalized to the amount of extracted protein and is reported as µmol/g protein. Protein was quantified using the bicinchoninic acid assay.

#### NMR analysis

Lyophilized polar extracts were reconstituted in D2O (> 99.9%, Cambridge Isotope Laboratories, MA) containing 17.5 nmol d6-2,2-dimethyl-2-silapentane-5-sulfonate (DSS) as internal standard were analyzed by 1D ^1^H (PRESAT) and ^1^H(^13^C) -HSQC NMR on a Bruker 16.45 Avance III spectrometer equipped with a 1.7 mm HCN triple resonance cryoprobe. 1D ^1^H spectra were acquired at 15°C with 512 transients, 16384 data points, 12 ppm spectral width, an acquisition time of 2 s and a 6 s recycle time with weak irradiation on the residual HOD signal during the relaxation delay. The raw fids were apodized with 1 Hz exponential line broadening and zero filled to 131072 points prior to Fourier transformation. 1D ^1^H(^13^C)HSQC spectra were recorded with an acquisition time of 0.25 s with adiabatic decoupling and recycle time of 2 s over a spectral width of 12 ppm, with, 1024 transients. The HSQC spectra were then apodized with unshifted Gaussian function and 4 Hz exponential line broadening and zero filled to 16384 data points before Fourier transformation. Metabolites were assigned by comparison with in-house^21^ and public NMR databases. Metabolites and their ^13^C isotopomers were quantified using the MesReNova software (Mestrelab, Santiago de Compostela, Spain) by peak deconvolution. The peak areas of metabolites obtained were converted into nmoles by calibration against the peak intensity of DSS (17.5 nmoles) at 0 ppm for ^1^H spectra. Peak areas for each spectrum were corrected for the number of protons, calibrated against DSS to determine the absolute amount, and then normalized to tissue residue weight to obtain metabolite content as µmol/g residue. The fractional enrichment of site specific ^13^C (F) was calculated from the ^1^H spectra as:

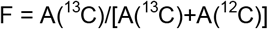

where A(^13^C) and A(^12^C) are the peak areas of the proton of interest directly attached to ^13^C and ^12^C, respectively.

### In vivo studies

#### Hyperinsulinemic-hypoglycemic clamps

The glucose clamp study in mice was performed at UMass Metabolic Disease Research Center as described previously^22^ with some modifications. Blood glucose levels were clamped around 60 mg/dl using continuous insulin and glucose infusions. At the end of the clamp study, 2-[1-^14^C]deoxy-d-glucose was injected into the mice and tissues of interest were collected 45 min. after the injection to determine tissue-specific insulin-stimulated glucose uptake.

#### Sympathetic nerve activity (SNA)

The adrenal and hepatic SNA were measured as described previously^23^. After anesthesia, the nerve innervating the liver or adrenal gland was identified, dissected free, and placed on a bipolar 36-gauge platinum-iridium electrode (A-M Systems; Carlsborg, WA). The electrode was connected to a high-impedance probe (HIP-511; Grass Instruments Co., Quincy, MA), and the nerve signal was amplified 10^5^ times with a Grass P5 AC pre-amplifier and filtered at low and high-frequency cutoffs of 100 Hz and 1000 Hz, respectively. This nerve signal was directed to a speaker system and to an oscilloscope (54501 A, Hewlett–Packard Co., Palo Alto, CA) for auditory and visual monitoring of the nerve activity. The signal was then directed to a resetting voltage integrator (B600C, University of Iowa Bioengineering) that sums the total voltage output in units of 1 V × s before resetting to zero and counting the number of spikes per second. The final neurograms were continuously routed to a MacLab analogue–digital converter (8 S, AD Instruments Castle Hill, New South Wales, Australia) for permanent recording and data analysis on a Macintosh computer. The nerve activity was measured for 30 min.

#### Blood glucose measurement

V-9302 (75 mg/kg bodyweight, dissolved in 5% DMSO / 95% sesame oil, %v/v, A1002473, Ambeed), dapagliflozin (10 mg/kg bodyweight, dissolved in PBS, A120381, Ambeed) and sotagliflozin (30 mg/kg bodyweight, dissolved in 5% DMSO / 95% sesame oil, %v/v, A225622, Ambeed) were administered intraperitoneally to different cohorts of non-fasted C57BL/6J mice (000664, Jackson Laboratory). The mouse tail-end was nicked using a razor blade and the tail was gently massaged to obtain a drop of blood. A glucometer (Alpha Trak 3, Zoetis) was then used per the manufacturer’s instructions to measure the blood glucose levels at different times after the administration of drugs.

To achieve kidney-specific knockout of *Mpi*, adult *Mpi*^*loxp/loxp*^ mice (T016292, GemPharmatech) were anesthetized, and the kidneys were exposed via a midline laparotomy. AAV9-CMV-Cre (105537, Addgene), or AAV9-CMV-GFP as a control (105530, Addgene), in a 50 µL volume, was delivered via retrograde renal pelvis injection using a 33-gauge needle. To maximize transduction efficiency and minimize systemic leakage, the renal artery and vein were temporarily occluded with micro-vascular clamps for 15 minutes during and after the injection. Three weeks after AAV administration, mice were treated with dapagliflozin, and blood glucose levels were monitored as described above.

*Mpi* knockout efficiency in the kidney was quantified by RT-qPCR using *Hprt* as the internal reference gene for normalization. The following primers were used: *Mpi: 5’-* CAGCACAGAGTCAAGATGACCC-3’ and 5’-GAAGGATGCTGGCAGAGTCTAG-3’; *Hprt: 5’-*CTGGTGAAAAGGACCTCTCGAAG-3’ and 5’-CCAGTTTCACTAATGACACAAACG-3’

### Molecular and cellular analyses

We used a mouse cytokine array (ARY028, R&D systems) to analyze serum of r*Glut2* KO mice and their littermate controls per the manufacturer’s instructions. We collected 50 µl serum from each mouse. Then, the serum of three mice in each group was pooled into 2ml microtubes separately for the control and experimental groups. The array membranes were developed using chemiluminescent detection reagents, imaged via ChemiDoc XRS+ (Bio-Rad Laboratories) and analyzed through Image Lab 6.1 software.

For multiomic gene analysis, we isolated nuclei from three flash-frozen mouse kidney samples per group and constructed single nucleus gene expression libraries and single nucleus transposase-accessible chromatin regions (ATAC) libraries on the 10x genomics chromium system using the 10x chromium multiome kit (Singulomics Corporation). Each library was sequenced with ∼200 million pair end reads (PE150) on Illumina NovaSeq. Using the 10x solution, we assayed for ATAC alongside gene expression (RNA-seq) for combined epigenomic and transcriptomic analysis from the same single nucleus. For each sample, we applied Seurat v5.1.0 and Signac v 1.13.0 to create a multiomic Seurat object with paired gene expression and DNA accessibility profiles for each sample. The sequencing reads were analyzed with mouse reference genome mm10 (2020-A) using the Cell Ranger ARC 2.0.2 count function. The transcription factor activity was predicted based on the presence of binding motifs within differential accessibility chromatin regions using chromVAR and the positional weight matrix (https://genome.ucsc.edu/goldenPath/help/hgRegMotifHelp.html) was obtained from the JASPAR2020 database (https://jaspar2020.genereg.net/). For each cell type, we identified differential accessibility cis-regulatory elements at ATAC-seq level and calculated marker genes at the RNA level. Additionally, we generated a gene activity matrix inferred from ATAC-seq by using the GeneActivity function using Signac.

Reduced-representation bisulfite sequencing (RRBS) was performed at Zymo Research, Irvine, CA. 500 ng of starting input genomic DNA was digested with 30 units of MspI (R0106, NEB). Fragments were ligated to pre-annealed adapters containing 5’-methyl-cytosine according to Illumina’s specified guidelines. Adaptor-ligated fragments ≥50 bp in size were recovered using the DNA Clean & Concentrator™-5 (D4003, Zymo Research). The fragments were then bisulfite-treated using the EZ DNA Methylation-Lightning™ Kit (D5030, Zymo Research). Preparative-scale PCR was performed and the resulting products were purified with DNA Clean & Concentrator™-5 (D4003, Zymo Research). Library quality control was performed on the Agilent 4200 TapeStation. Libraries were sequenced on an Illumina NovaSeq X platform (150 bp PE reads). The sequence reads from RRBS libraries (non-directional) were identified using standard Illumina base calling software. To process RRBS data, the trimmed reads were aligned to the mouse genome assembly using Bismark. Methylation ratios for each cytosine in CpG context were called using MethylDackel. The methylation level of each sampled cytosine was estimated as the number of reads reporting a C, divided by the total number of reads reporting a C or T. Read depths per cytosine in the genome as well as in different genomic regions (e.g. gene body, promoter, CpG island, etc. based on available annotations) were calculated and tabulated using scripts.

### Statistics

In vivo data are shown as mean□± SEM. Results were analyzed by two-tailed Student’s unpaired t-test, multiple comparison t-test, or repeated measures two-way ANOVA followed by a Holm-Sidak or Bonferroni post hoc multiple comparison test when appropriate. All analyses were performed using Prism version 10.2.3 (GraphPad, USA) and differences were considered statistically significant at p<0.05.

For multiomic single nucleus RNA and ATAC sequencing, the analysis of each cell type was carried out using the Wilcoxon Rank Sum Test, implemented through the Seurat FindAllMarkers function. The differentially expressed genes and accessibility were considered statistically significant at false discovery rate (FDR) adjusted p<0.05. The pathway analysis for cross-condition comparisons involves evaluating overexpressed and underexpressed genes within each cell type using Gene Set Enrichment Analysis (GSEA). GSEA considers the magnitude of differential expression for each gene by applying gene-specific weights. The pathways selected for analysis are obtained from the Molecular Signatures Database through the R package msigdbr.

To compare RRBS results between the groups, a statistical analysis was performed to identify, annotate, and visualize differential methylated sites and regions using differential shrinkage sequencing (DSS) package (software Bioconductor 3.22). Differentially methylated cytosines (DMCs) and differentially methylated regions (DMRs) were detected using DSS, which by default conducted the Wald test and the Benjamini-Hochberg P-value adjustment. Significant DMCs and DMRs have FDR ≤ 0.05 and absolute methylation difference ≥ 0.1.

## Results

### Renal glycosuria induces metabolic rewiring in the kidney

To understand how systemic glucose homeostasis is maintained during chronic renal glycosuria, we examined metabolic adaptations in the kidney using untargeted metabolomics in r*Glut2* KO mice. We observed that mannose levels were about 18-fold higher in the KO mice compared with their control littermates (figure 1A and line 99, supplemental table 1). The KO mice also had high levels of glutamine, tyrosine and glycine (figure 1A; lines 145, 25, and 140, respectively, supplemental table 1), suggesting altered amino acid handling, potentially including increased protein catabolism or reduced amino acid utilization. In contrast, monosaccharides such as galactose and fructose were decreased (lines 153 and 159, respectively, supplemental table 1) in the KO mice. This metabolic profile resembles adaptations observed during glucose deprivation and suggests that alternative carbon sources, including mannose and glutamine, may contribute to maintaining glucose homeostasis during renal glycosuria. We focused on examining these two metabolites for subsequent analyses because mannose has been suggested to support energy homeostasis during high metabolic demands^24-26^ and glutamine is a major substrate for gluconeogenesis in the kidney^27-29^.

**Figure 1.**
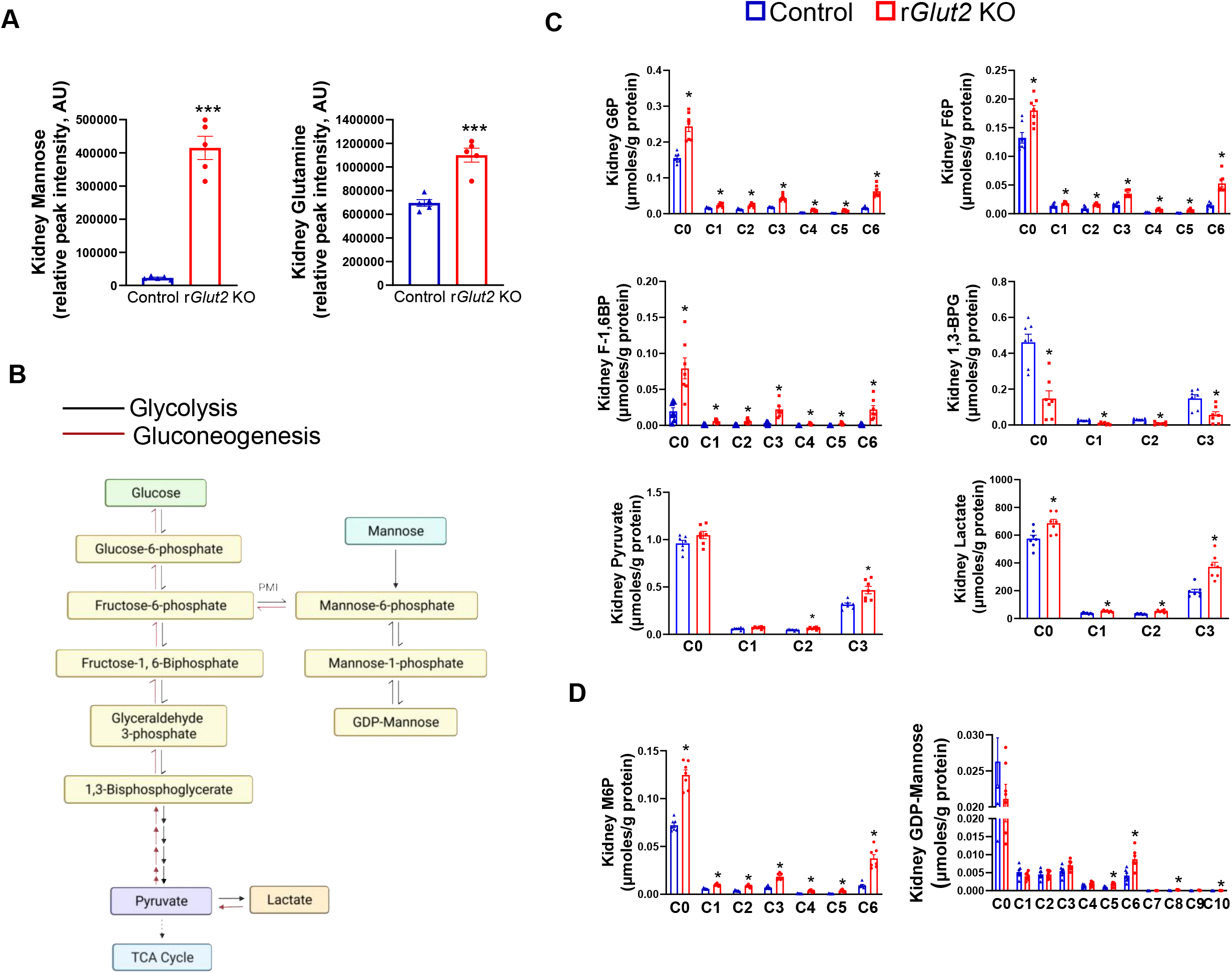
Untargeted metabolomics and stable isotope-resolved metabolomics analysis (by Ion Chromatography-Ultra-High Resolution Fourier Transform Mass Spectrometry) of kidney glucose and mannose metabolism in 14-week-old male renal glucose transporter 2 knockout (r*Glut2* KO) mice and their control littermates 60 minutes after ^13^C_6_ glucose administration. **(A)** Untargeted metabolomics analysis showing increased mannose and glutamine levels in r*Glut2* KO, N=5, ***p<0.0001, two-tailed unpaired t-test. **(B)** Schematic of glycolysis, gluconeogenesis, and mannose metabolism. **(C)** ^13^C-glucose labeling of glucose 6-phosphate (G6P), fructose 6-phosphate (F6P), fructose 1,6, bisphosphate (F1,6BP), and lactate was increased, whereas labeling of 1,3-bisphosphoglycerate (1,3-BPG) and pyruvate was decreased in r*Glut2* KO kidneys. This discontinuity suggests bidirectional carbon flux consistent with glucose cycling and reversible steps within the upper glycolytic pathway. The X-axis represents the isotopologue distribution (number of ^13^C atoms) of each metabolite. **(D)** ^13^C-glucose incorporation into mannose 6-phosphate (M6P) and GDP-mannose was increased in r*Glut2* KO kidneys, indicating enhanced routing of hexose carbons into the mannose/M6P pathway during glucose loss. N=7, *p<0.05, multiple comparison unpaired t-test corrected using Holm-Sidak method. PMI, phosphomannose isomerase (also known as mannose 6-phosphate isomerase, MPI)

To comprehensively investigate the role of glucose and mannose metabolism in energy homeostasis during renal glycosuria, we used a stable isotope tracing technique. We administered ^13^C_6_ glucose (oral gavage, 60 mg/mouse) to mice and tracked the incorporation of ^13^C into downstream metabolites 60 min. after the labeled glucose administration. In the kidneys of r*Glut2* KO mice, the ^13^C incorporation was increased in proximal glycolytic intermediates, including glucose-6-phosphate (G6P) and fructose-6-phosphate (F6P), as well as in downstream metabolites such as pyruvate and lactate (figure 1B,C). At the same time, enrichment of the intermediate metabolite 1,3-bisphosphoglycerate (1,3-BPG) showed no proportional increase relative to upstream intermediates. This discontinuity in labeling across glycolysis suggests bidirectional carbon flux, including reversible upper-glycolytic reactions and/or enhanced glucose cycling within the pathway. ^13^C-glucose incorporation into mannose metabolites was increased in the kidney (figure 1D), suggesting that during renal glycosuria glucose appears to enter mannose metabolism pathways via the enzyme mannose 6-phosphate (M6P) isomerase (MPI, also known as phosphomannose isomerase (PMI)) as described in Figure 1B. This finding indicates increased routing of hexose carbons through the mannose metabolism pathway during glucose loss.

To directly determine the role of mannose in mediating the glucose-sparing or - producing effects observed during renal glycosuria, we administered ^13^C_6_ mannose (oral gavage, 60 mg/mouse) to r*Glut2* KO mice and their control littermates. We then analyzed glucose as well as mannose metabolism pathways 60 min. after the labeled mannose administration. We observed that incorporation of ^13^C-mannose into hexose-phosphate and upper-glycolytic intermediates, including M6P, was higher in the kidneys of the KO mice compared with their littermate controls (figure 2). The contribution of ^13^C-mannose to kidney GDP-mannose is reduced (figure 2B) in the KO mice, indicating the accumulation of M6P for gluconeogenic or upstream metabolic pathways.

**Figure 2.**
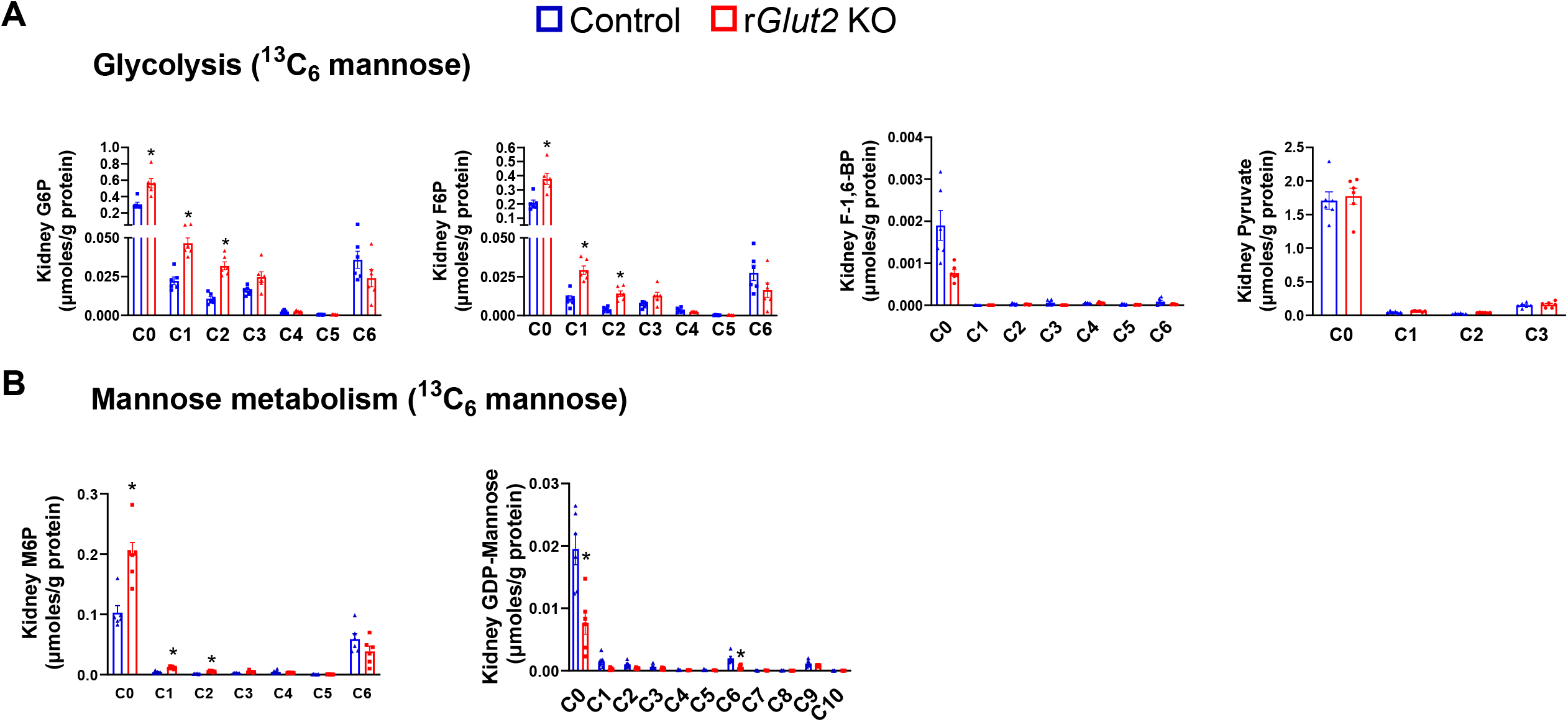
Stable isotope-resolved metabolomics analysis (by Ion Chromatography-Ultra-High Resolution Fourier Transform Mass Spectrometry) of kidney glycolysis and mannose metabolism in 14-week-old male renal glucose transporter 2 knockout (r*Glut2* KO) mice and their control littermates 60 minutes after ^13^C_6_ mannose administration. ^13^C-mannose incorporation into upper-glycolytic and hexose-phosphate intermediates **(A)**, and into mannose-derived intermediates **(B)** was increased (except fructose 1,6-bisphosphate (F1,6BP) and GDP-mannose, which were not changed or decreased) in r*Glut2* KO kidneys. These patterns indicate increased mannose entry into the hexose-phosphate pool and diversion toward upstream metabolic pathways during glycosuria. The X-axis represents the isotopologue distribution (number of ^13^C atoms) of each metabolite. N=7, *p<0.05, multiple comparison unpaired t-test corrected using Holm-Sidak method. G6P, glucose 6-phosphate; F6P, fructose 6-phosphate.

NMR analysis of ^13^C_6_ mannose tracing showed fractional enrichment of ^13^C-labeled glucose in the urine of r*Glut2* KO mice (∼8.5 ±3%, mean ±SD) but not in the control group (glucose was undetectable in the control mice). This finding indicates conversion of mannose to glucose in KO mice through the pathways delineated in figure 1B. In addition, ^13^C-mannose-derived glucose and lactate (a gluconeogenic substrate) were increased in the kidneys of KO mice compared with control littermates (supplemental figure 1), further supporting a contribution of mannose to renal glucose production during glycosuria. Together, the ^13^C-mannose tracing and urinary ^13^C-glucose enrichment demonstrate net mannose-to-glucose conversion in vivo, supporting substrate-level glucose production through the hexose-phosphate pathway during renal glycosuria.

In the skeletal (soleus) muscle, ^13^C-glucose incorporation into glycolytic and mannose-related intermediates was reduced during glycosuria (figure 3A, B), indicating reduced glycolytic flux and/or diminished glucose utilization. By contrast, ^13^C-mannose labeling of G6P, F6P, and downstream mannose-related intermediates increased (figure 3C, D), indicating greater mannose uptake and entry into the hexose-phosphate pool. However, the decrease in F-1,6-BP together with unchanged pyruvate enrichment shows that mannose carbons do not progress through the phosphofructokinase step into lower glycolysis. Instead, mannose is largely confined to the upper glycolytic/hexose-phosphate network. Thus, during renal glycosuria, skeletal muscle shifts substrate use toward mannose while limiting glucose-derived glycolysis.

**Figure 3.**
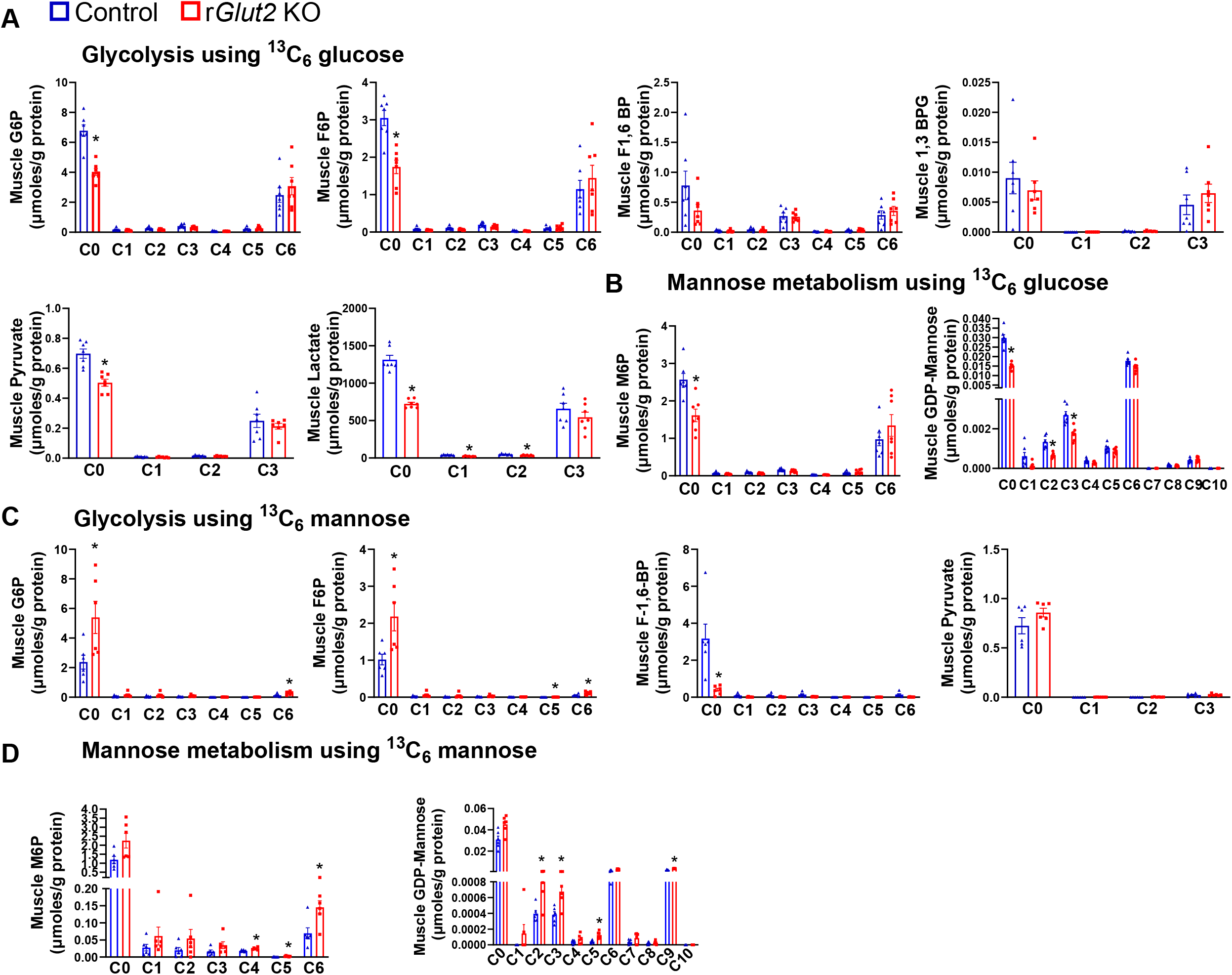
Stable isotope-resolved metabolomics analysis (by Ion Chromatography-Ultra-High Resolution Fourier Transform Mass Spectrometry) of soleus muscle glucose and mannose metabolism in 14-week-old male renal glucose transporter 2 knockout (r*Glut2* KO) mice and their control littermates 60 minutes after ^13^C_6_-glucose or ^13^C_6_-mannose administration. **(A)** ^13^C-glucose labeling of metabolites such as glucose 6-phosphate (G6P), fructose 6-phosphate (F6P), fructose 1,6, bisphosphate (F1,6BP) and pyruvate was decreased in r*Glut2* KO mice, indicating reduced muscle glycolytic flux. The X-axis represents the isotopologue distribution (number of ^13^C atoms) of each metabolite. **(B)** ^13^C-glucose incorporation into muscle mannose-6-phosphate (M6P) and GDP-mannose was reduced in r*Glut2* KO mice, indicating decreased glucose-mediated mannose metabolism. Conversely, ^13^C-mannose labeling of hexose-phosphate and glycolytic intermediates **(C)** and mannose metabolites **(D)** was increased in r*Glut2* KO mice, except for F1,6BP, which was decreased. These patterns indicate that mannose increasingly enters upper glycolysis in muscle during glycosuria while glucose utilization is suppressed. N=7, *p<0.05, multiple comparison unpaired t-test corrected using Holm-Sidak method. 1,3-BPG, 1,3-bisphosphoglycerate.

To determine how these glycolytic shifts affect downstream oxidative metabolism, we examined ^13^C incorporation into TCA-cycle intermediates. In the kidney, incorporation of both ^13^C-glucose and ^13^C-mannose into TCA-cycle metabolites increased during renal glycosuria (Figure 4). Conversely, in skeletal muscle, ^13^C incorporation into TCA-cycle intermediates increased only after ^13^C-mannose administration (Figure 5). The skeletal muscle α-ketoglutarate was increased under both tracer conditions (Figures 4 and 5), likely reflecting anaplerotic inputs from amino acids such as glutamine and glutamate. In the kidney, increased labeling of upstream TCA intermediates and elevated α-ketoglutarate are similarly consistent with enhanced glutamine-supported anaplerosis during renal glycosuria. These TCA-cycle findings align with the glycolysis results described above, indicating that skeletal muscle relies on mannose for oxidative metabolism during glycosuria. Likewise, the combined findings from Figures 1D and 4B suggest that renal glucose is routed through the mannose/M6P pathway to support TCA-cycle activity during glycosuria.

**Figure 4.**
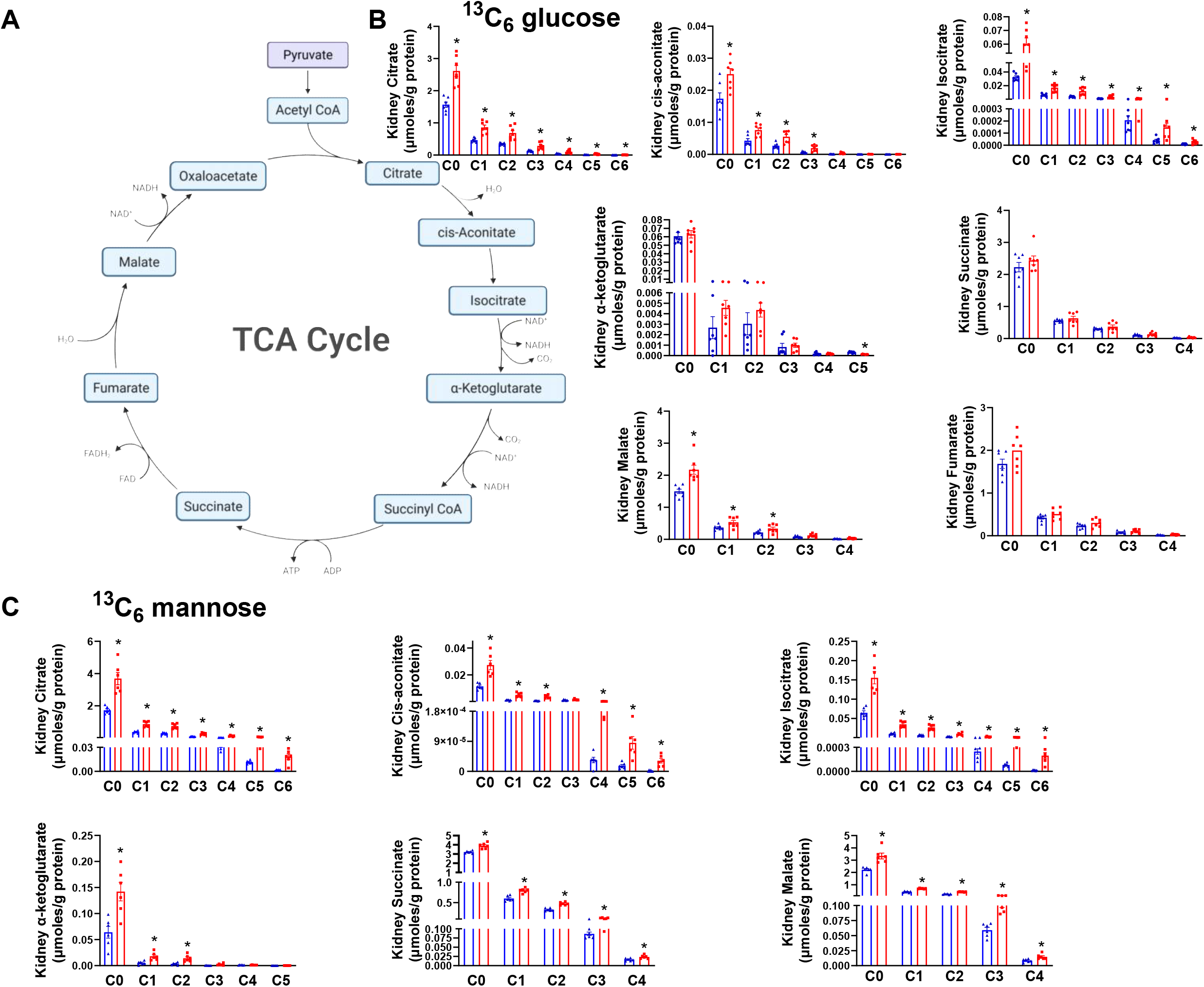
Stable isotope-resolved metabolomics analysis (by Ion Chromatography-Ultra-High Resolution Fourier Transform Mass Spectrometry) of kidney tricarboxylic acid (TCA) cycle metabolites in 14-week-old male renal glucose transporter 2 knockout (r*Glut2* KO) mice and their control littermates 60 minutes after ^13^C_6_ glucose or ^13^C_6_ mannose administration. **(A)** Schematic of the TCA cycle. **(B, C)** ^13^C-glucose and ^13^C-mannose incorporation into citrate, isocitrate, cis-aconitate, and α-ketoglutarate was increased in r*Glut2* KO kidneys. Succinate ^13^C labeling increased only after ^13^C-mannose administration, whereas fumarate showed little or no change under either condition. These findings indicate renal metabolic reprogramming that enhances substrate-level carbon entry into the TCA cycle. The X-axis represents the isotopologue distribution (number of ^13^C atoms) of each metabolite. N=7, *p<0.05, multiple comparison unpaired t-test corrected using Holm-Sidak method.

**Figure 5.**
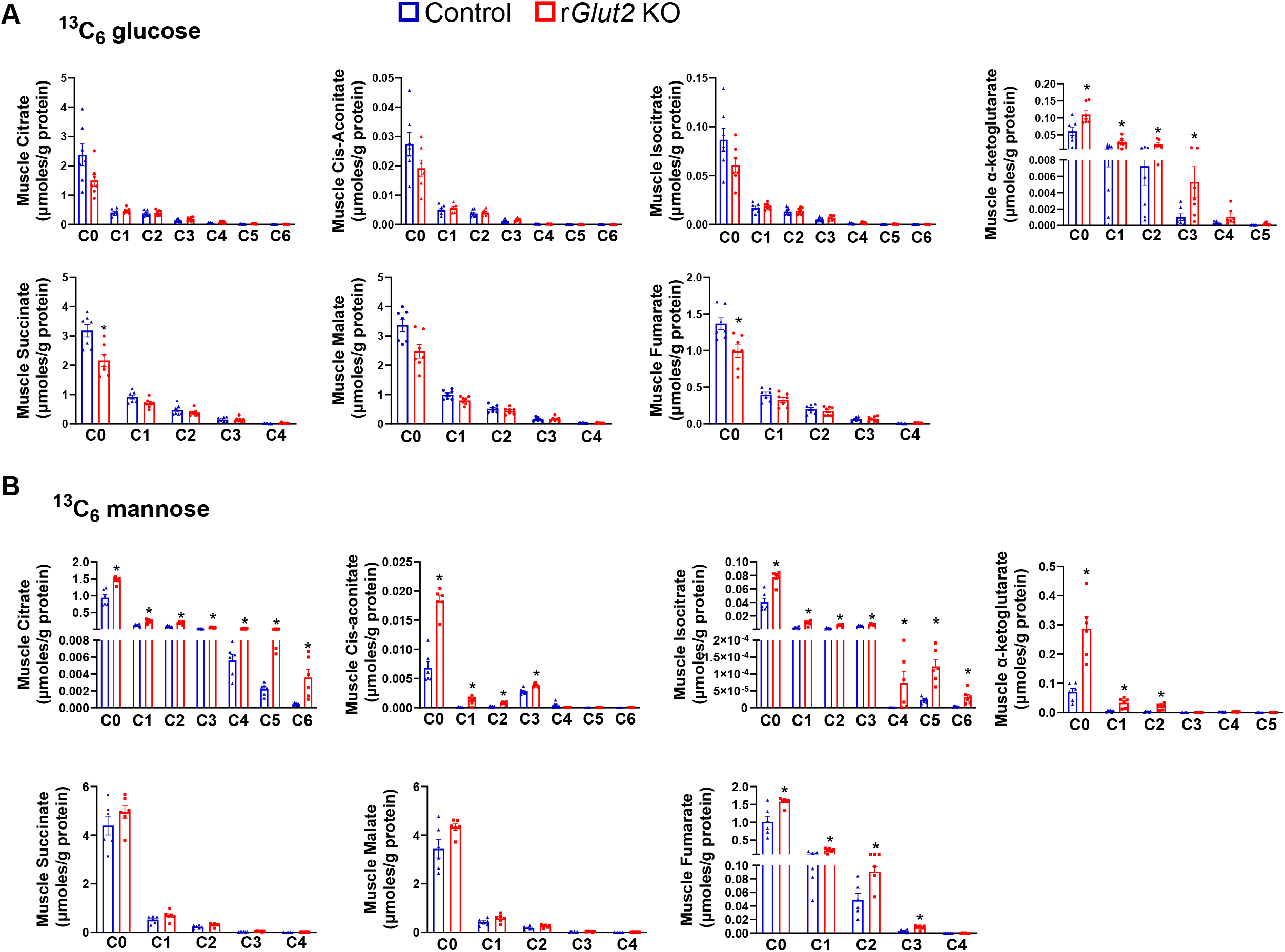
Stable isotope-resolved metabolomics analysis (by Ion Chromatography-Ultra-High Resolution Fourier Transform Mass Spectrometry) of soleus muscle tricarboxylic acid (TCA) cycle metabolites in 14-week-old male renal glucose transporter 2 knockout (r*Glut2* KO) mice and their control littermates 60 minutes after ^13^C_6_ glucose or ^13^C_6_ mannose administration. **(A)** ^13^C-glucose labeling of TCA-cycle intermediates was reduced or unchanged in r*Glut2* KO mice, except for α-ketoglutarate, which was increased. **(B)** In contrast, ^13^C-mannose incorporation into TCA-cycle intermediates was increased (except for succinate and malate, which were unchanged), indicating that mannose increasingly fuels muscle oxidative metabolism during renal glycosuria and contributes to skeletal-muscle glucose sparing. The X-axis represents the isotopologue distribution (number of ^13^C atoms) of each metabolite. N=7, *p<0.05, multiple comparison unpaired t-test corrected using Holm-Sidak method.

Unexpectedly, the liver did not show major changes in the glucose and mannose metabolism (supplemental figures 2 and 3) at the timepoint used in this study. The liver plays a key role in gluconeogenesis during fasting, including in r*Glut2* KO mice, as reported previously^12^. However, because ^13^C_6_ glucose was administered in this study, the liver may have perceived a fed state at the time of metabolic analysis, thereby attenuating detectable changes in glucose metabolism. By contrast, the kidney exhibited marked metabolic alterations in response to glycosuria.

Collectively, the IC-UHR-FTMS and NMR analyses indicate that mannose plays a key role in maintaining glucose homeostasis during renal glycosuria by supplying carbon to the hexose-phosphate pool for substrate-level glucose production in the kidney and by supporting oxidative metabolism in skeletal muscle.

### Renal glycosuria induces systemic glucose-sparing physiology

To determine whether renal glycosuria triggers systemic metabolic adaptations that facilitate glucose sparing, we examined whole-body glucose metabolism and autonomic activity. Hyperinsulinemic–hypoglycemic clamps (figure 6A) showed that r*Glut2* KO mice maintained euglycemia without requiring exogenous glucose infusion (figure 6B). The KO mice show normal hepatic insulin sensitivity (Figures 6C, D); however, the whole-body glycolysis is reduced (figure 6E). While the glucose uptake is unaffected in the epididymal white adipose tissue (Figure 6F), we observed a decrease in glucose utilization (insulin resistance) in the gastrocnemius muscle (Figure 6G) and brain (figure 6H) of the KO mice. The tissue-specific insulin resistance may explain why glucose metabolism was reduced (figure 3A, B) and mannose metabolism was higher (figure 3C, D, and figure 5B) in the skeletal muscle of r*Glut2* KO mice compared with the control group. These systemic glucose sparing adaptations in r*Glut2* KO mice may clarify how glucose is made available for necessary biological activities including the counterregulatory response.

**Figure 6.**
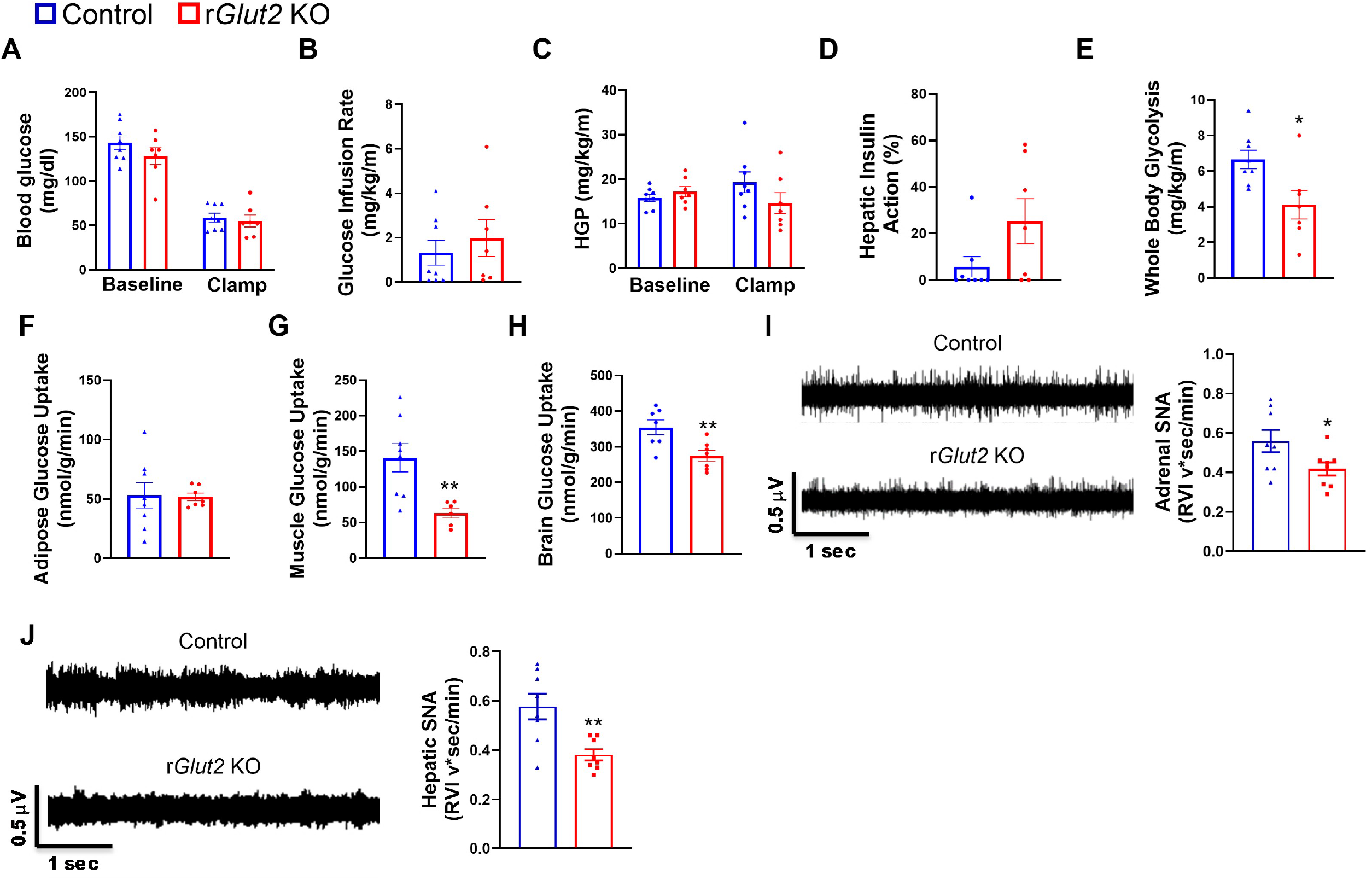
Hyperinsulinemic-hypoglycemic clamp and sympathetic nerve activity (SNA) measurements in 12-week-old male renal glucose transporter 2 knockout (r*Glut2* KO) mice and their control littermates. **(A)** Blood glucose levels during baseline and hypoglycemic clamp. **(B)** Glucose infusion rate (GIR), which was unchanged between groups. **(C–D)** Hepatic glucose production (HGP) and hepatic insulin action, showing preserved hepatic insulin sensitivity in r*Glut2* KO mice. **(E)** Whole-body glycolysis was reduced in r*Glut2* KO mice. **(F–H)** Tissue-specific glucose uptake: unchanged in epididymal white adipose tissue **(F)**, reduced in gastrocnemius muscle **(G)**, and reduced in the brain **(H)**, consistent with tissue-specific insulin resistance. **(I–J)** Adrenal **(I)** and hepatic **(J)** SNA were reduced in r*Glut2* KO mice; representative SNA tracings are shown on the left and quantification on the right. N=7 or 8, *p<0.05, **p<0.01, two-tailed unpaired t-test.

Because sympathetic nerve activity contributes to counterregulation during hypoglycemia, we also measured hepatic and adrenal sympathetic output. Both signals were markedly suppressed in r*Glut2* KO mice (figure 6I, J), consistent with a systemic energy-conserving phenotype. These findings support the view that renal glycosuria elicits physiological responses that reduce whole-body glucose utilization and complement the substrate-level rewiring observed in this study.

### Mannose and Glutamine contribute to glucose compensation during renal glycosuria

Given that renal glycosuria increased mannose and glutamine levels as described above, we next determined whether inhibiting mannose production or blocking glutamine metabolism would impair glucose homeostasis during renal glycosuria.

We administered V-9302^30^ - a glutamine transporter ASCT2 (encoded by *Slc1a5* gene) inhibitor – to the mice to determine whether glutamine metabolism contributes to glucose compensation in renal glycosuria. We monitored blood glucose levels at different times following the V-9302 administration (figure 7A). Remarkably, after the initial increase in blood glucose levels, the glucose concentration in the r*Glut2* KO mice decreased to potentially life-threatening hypoglycemia compared with the control mice. The KO mice had to be given a glucose bolus (60 mg/mouse, intraperitoneally) to restore normal conditions. The initial increase in the blood glucose levels was likely due to a concomitant increase in circulating glutamine levels, which eventually decrease with time, as reported previously^30^. Next, we administered V-9302 (once a day) or its vehicle to C57BL6/J wildtype mice (000664, the Jackson laboratory) that were previously treated with the SGLT2 inhibitor dapagliflozin (twice a day), and monitored their blood glucose levels. V-9302 was administered one hour after the second dose of dapagliflozin for two days. The V-9302 and dapagliflozin treated mice showed hypoglycemia compared with the mice that received the vehicle and dapagliflozin (figure 7B), thereby supporting the findings from V-9302-treated r*Glut2* KO mice.

**Figure 7.**
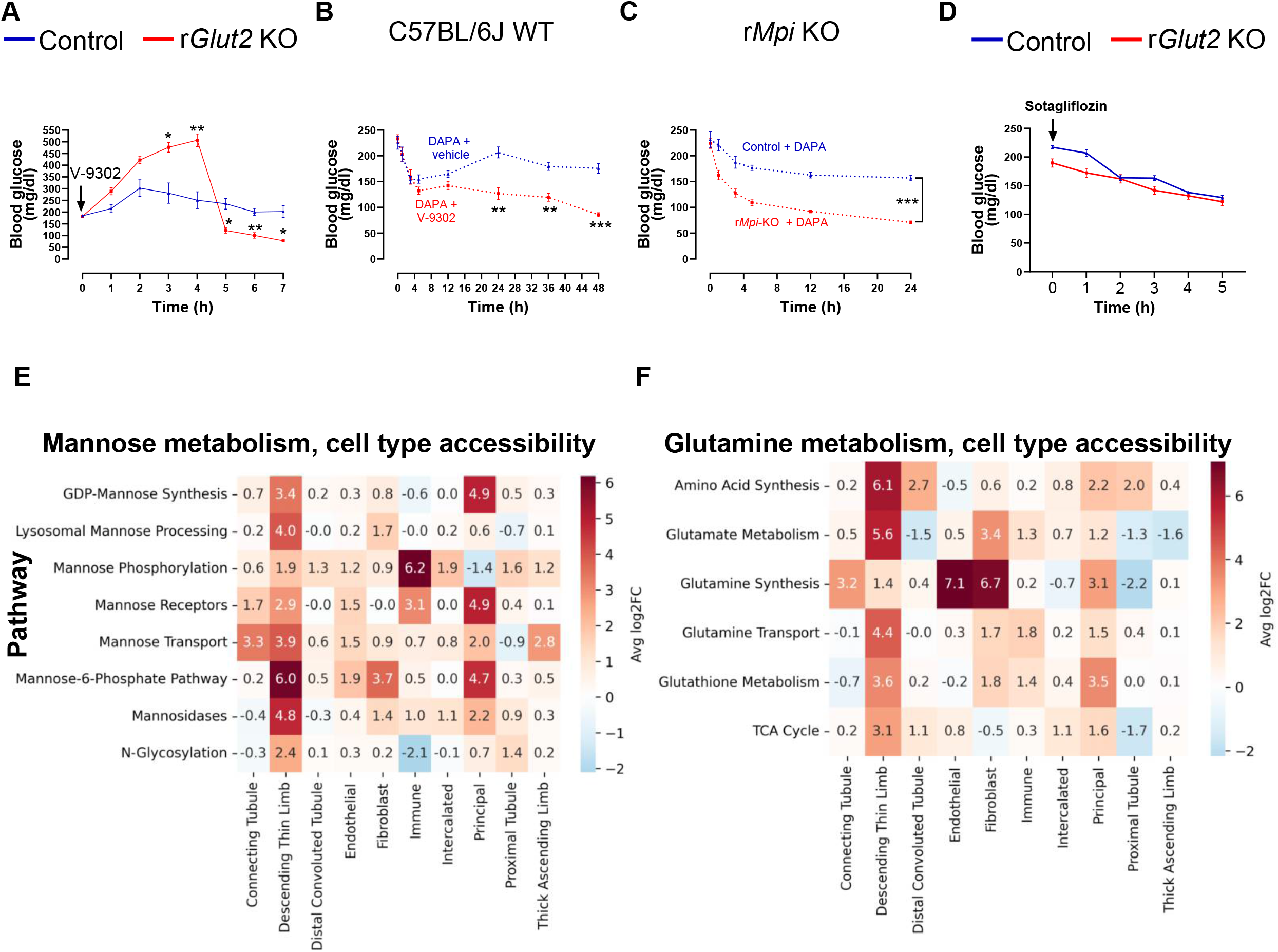
Effects of V-9302 (an inhibitor of glutamine transport/metabolism), dapagliflozin (DAPA; SGLT2 inhibitor), and sotagliflozin (dual SGLT1/SGLT2 inhibitor) on blood glucose levels, and chromatin accessibility of mannose and glutamine metabolic pathways. **(A)** In 26-week-old male r*Glut2* KO mice, V-9302 (75□mg/kg) induced hypoglycemia following an initial rise in blood glucose. **(B)** In 9-week-old WT mice treated with DAPA (10□mg/kg, twice daily), V-9302 (75□mg/kg, once daily; 1□h after the second DAPA dose) produced a similar hypoglycemic effect. **(C)** DAPA induced hypoglycemia in 20-week-old AAV-mediated renal *Mpi* knockout (r*Mpi* KO) mice. **(D)** Sotagliflozin (30□mg/kg) reduced blood glucose similarly in control and r*Glut2* KO mice. **(E–F)** ATAC-seq analysis of kidneys from 12-week-old male r*Glut2* KO and control mice showed increased chromatin accessibility in mannose **(E)** and glutamine **(F)** metabolic pathways, consistent with enhanced substrate availability and reduced catabolic activity. N□=□6. *p□<□0.05, **p□<□0.01; repeated-measures two-way ANOVA with Bonferroni post hoc correction.

To block renal mannose production, we deleted *Mpi* in the kidney of *Mpi*^flox/flox^ mice by delivering AAV9-CMV-Cre via retrograde renal pelvis injection; AAV9-CMV-GFP was used as the control. Three weeks after AAV administration, mice were treated with dapagliflozin (twice a day) and blood glucose levels were monitored as described above. *Mpi* deficient mice exhibited hypoglycemia compared with their control group (figure 7C). Renal *Mpi* deletion was confirmed by RT-qPCR (6.3 ± 2% of control expression; controls: 100 ± 31%; n = 7, mean ±SEM) at the end of the study. These findings indicate that mannose contributes to maintaining glucose homeostasis during renal glycosuria.

Because kidney SGLT1 is assumed to compensate for SGLT2 inhibition^5^, a dual SGLT1 and SGLT2 inhibitor could improve the efficiency of SGLT2 inhibition alone. To test this hypothesis, we monitored blood glucose levels in r*Glut2* KO mice after administering sotagliflozin (dual SGLT1 and SGLT2 inhibitor). The drug reduced blood glucose levels in both the KO mice and their control littermates to a similar extent (figure 7D). This observation indicates that glucose reabsorption by kidney SGLT1 may not fully explain how r*Glut2* KO mice maintain normal blood glucose levels in renal glycosuria. We have previously shown that kidney SGLT2 is reduced, but SGLT1 is unaffected, in r*Glut2* KO mice^2^. Altogether, these data indicate that mannose utilization and glutamine metabolism play an important role in supporting glucose compensation during renal glycosuria.

### Kidney molecular adaptations and systemic stress during renal glycosuria

To determine whether molecular changes in the kidney contribute to the metabolic adaptations observed in vivo, we analyzed kidney tissue using single-nucleus RNA-seq, ATAC-seq, and reduced-representation bisulfite sequencing (RRBS). Both RNA and chromatin accessibility data (figure 7E, supplemental tables 2 and 3) showed coordinated downregulation of genes involved in glutamine catabolism (with reduced chromatin accessibility in ATAC-seq) and upregulation of genes involved in glutamine transport, including *Slc1a5* (higher chromatin accessibility in ATAC-seq). Similarly, genes involved in mannose metabolism, including *Mpi*, exhibited increased accessibility and the mannose transporter gene *Slc5a10* was upregulated in the KO mice (figure 7F, supplemental tables 2 & 3). *Mpi* encodes mannose 6-phosphate isomerase, which interconverts mannose-6-phosphate and fructose-6-phosphate, supporting the glucose and mannose shunt identified in the ^13^C-tracing studies. RRBS analysis (supplemental table 4) further showed hypomethylation of *Slc1a5* and hypermethylation of genes involved in glutamine catabolism such as *Got2*, consistent with enhanced glutamine uptake and reduced oxidative glutamine catabolism. These molecular adaptations may increase intracellular glutamine availability for gluconeogenic substrate supply rather than for conventional mitochondrial glutamine oxidation, aligning with the substrate-level rewiring suggested by the ^13^C-tracing experiments.

To further assess systemic signals associated with glucose sparing, we measured circulating cytokines using a microarray dot plot. Several markers of metabolic stress and immune activation, including CCL6, GDF15, endostatin, and IGFBP1, were elevated in r*Glut2* KO mice, whereas cytokines associated with immunosuppressive signaling such as VEGF, RAGE, and PDGF-BB were reduced (supplemental figure 4). These results extend our previous findings that renal glycosuria elicits an acute phase response including activation of the immune system^12^. Collectively, these changes indicate systemic metabolic stress and immune activation, which may contribute to the kidney’s adaptive metabolic rewiring in response to glucose loss.

To evaluate whether classical gluconeogenic genes were altered, we examined *Pck1* and *G6pc* in our transcriptomic and ATAC-seq datasets (supplemental tables 2 & 3). In the RNA-seq analysis, *Pck*1 was not detectable in any kidney cell types and *G6pc* was reduced in proximal tubule cells of r*Glut2* KO mice (log2FC -1.90; p adj. = 3.4×10^−2^□). ATAC-seq analysis aligned with these findings: neither locus showed increased chromatin accessibility under the experimental condition. Instead, both *Pck1* and *G6pc* peaks showed slightly higher accessibility in the control group (fold changes 0.60 and 0.67). These data indicate that changes in *Pck1* or *G6pc* transcription or accessibility are unlikely to explain the substrate-level glucose production observed in our ^13^C-tracing experiments. Therefore, our findings support the conclusions reported previously that the kidney adjusts its gluconeogenic activity largely through changes in substrate delivery, rather than through major transcriptional upregulation of *Pck1* or *G6pc*^27,31^. Notably, metabolic flux studies demonstrate that *Pck1* (which encodes PEPCK-C) expression does not necessarily predict gluconeogenic flux^32^.

Overall, the molecular analyses suggest that renal glycosuria triggers a coordinated metabolic reprogramming - favoring mannose and glutamine accumulation, reduced catabolism, and increased substrate availability to support glucose homeostasis. These findings align with the in vivo metabolic flux studies and indicate that substrate-level regulation is a key mechanism of glucose compensation during renal glycosuria.

## Discussion

The glucose lowering efficacy of SGLT2 inhibitors is attenuated by compensatory mechanisms that maintain circulating glucose levels, including endogenous glucose production and possible contributions from residual SGLT1-mediated reabsorption^5,13^. Prior work indicates that SGLT2 inhibition activates metabolic programs resembling starvation, promoting alternative energy use and conservation in models of type 2 diabetes^33,34^. Similarly, mice constitutively lacking *Glut2* in kidney proximal tubule cells show energy-conserving phenotype in response to renal glycosuria ^35^. However, the metabolic pathways that sustain glucose homeostasis during chronic renal glucose loss remain unclear.

Here, we investigated the metabolic pathways and associated molecular mechanisms that facilitate glucose compensation in renal glycosuria. To reduce potential confounding developmental effects associated with constitutive knockout models, we used a tamoxifen-inducible r*Glut2* KO mouse model^2^. Following renal *Glut2* deletion, these mice develop marked glycosuria accompanied by secondary loss of *Slc5a2* expression, resulting in sustained renal glucose wasting. This model therefore provides a genetic system to examine metabolic adaptations that support systemic glucose homeostasis during chronic renal glycosuria in adult mice. Metabolic profiling and stable isotope tracing revealed coordinated adaptations in mannose and glutamine metabolism that support glucose compensation.

Mannose metabolism has been described as an auxiliary energy pathway activated under glucose scarcity or high energy demand ^24-26^. Our present study shows that mannose metabolism is upregulated during renal glycosuria, and contributes to upper-glycolytic intermediates and to substrate-level glucose production via the hexose-phosphate pathway, thereby helping sustain glucose homeostasis. Elevated mannose may also contribute to tissue-specific insulin resistance, consistent with prior associations^36^ and with the reduced muscle glucose utilization observed in r*Glut2* KO mice. Together, these adaptations position mannose as a key substrate supporting both renal and systemic glucose sparing during glycosuria.

Glutamine is a major anaplerotic substrate that supports energy metabolism and precursor supply for glucose formation under conditions of metabolic stress^27-29^. In r*Glut2* KO mice, renal glutamine levels were elevated, and inhibition of glutamine transport with V-9302 caused hypoglycemia in both r*Glut2* KO and dapagliflozin-treated mice. These findings indicate that glutamine availability is required for maintaining glucose homeostasis during glycosuria. In addition, the increase in renal α-ketoglutarate during glycosuria is consistent with enhanced amino-acid–supported anaplerosis, as glutamine is well-established to be converted to glutamate and then to α-ketoglutarate for entry into the TCA cycle^28,37^. Together, these data identify glutamine metabolism as a critical component of the substrate-level adaptations that sustain glucose compensation during renal glucose loss.

Hormonal mediators of compensatory glucose production during glycosuria remain debated. Although early studies implicated glucagon secretion^38^, subsequent work^39^ – including ours^2^ - found no clear evidence for elevated glucagon following renal *Glut2* deletion or SGLT2 inhibition. Instead, glycosuric mice exhibit increased corticosterone and markers of metabolic stress^12^. In the present study, our data indicate that substrate-level metabolic adaptations involving mannose and glutamine play a central role in maintaining glucose homeostasis during glycosuria.

Importantly, SGLT2 inhibitors lower glucose without causing hypoglycemia in type 2 diabetes - likely because the metabolic pathways identified here maintain glucose homeostasis despite urinary glucose loss. These pathways provide a mechanistic basis for the preserved euglycemia observed during glycosuria and may help inform strategies to optimize the glucose-lowering efficacy of SGLT2 inhibitors while maintaining metabolic safety. Future work should define the kidney-derived signals that communicate renal glycosuria to the liver, muscle, and brain to coordinate systemic glucose sparing.

### Limitations

All experiments were conducted in non-diabetic mice with normal body weight and glucose levels to define physiological compensation to renal glycosuria. Whether mannose and glutamine metabolism make similar contributions in the context of obesity or diabetes remains to be determined. Although mannose is predominantly transported by SGLT4^40^ and SGLT5^41^, it is possible that lack of renal GLUT2 and SGLT2 may have indirectly altered mannose transport and metabolism in r*Glut2* KO mice. Similarly, SGLT4 and SGLT5 may contribute to glucose reabsorption in renal glycosuria as these transporters exhibit low affinity for glucose. Use of specific mannose transport inhibitors or kidney-specific *Mpi* KO models will further help clarify these possibilities. Additional studies are also needed to identify the renal cell types responsible for mannose production and to define the signals that induce mannose and glutamine metabolism during renal glycosuria. V-9302 inhibits additional amino acid transporters beyond ASCT2, so its effects may extend beyond glutamine transport^30^. Finally, we did not directly quantify contributions from individual gluconeogenic amino acids, lactate, or glycogenolysis to endogenous glucose production.

## Conclusion

Mannose and glutamine metabolism play key roles in supporting glucose compensation during renal glycosuria. These findings underscore kidney-driven metabolic adaptations that maintain systemic glucose homeostasis and provide a mechanistic framework that may inform strategies to optimize the glucose-lowering efficacy of SGLT2 inhibitors.

## Data Sharing Statement

The RNA, ATAC, and RRB sequencing data supporting the findings of this study are available in the supplementary data files 2, 3, and 4. The summary statistics are described and available in Methods. The software and code used in the analyses are cited in Methods.

## Acknowledgements

We thank Jason Kim, UMass Chan Medical School, for help with the clamp study, and Dalbir Kaur Chhabra for assistance with the bioinformatics analyses.

## Author contribution

N.R. and M.O. performed experiments, contributed to data analysis and manuscript editing; M.J.H., N.H. and H.Q. helped with tissue collection, mouse colony management including genotyping and preparing figures; T.F. and P.R.C performed the cytokine array and associated analyses; P.L., M.M.Y.K, and T.A.C performed the ^13^C glucose and mannose study including data analyses and manuscript editing; D.A.M. and K.R. performed the sympathetic nerve measurements and contributed to manuscript editing; K.H.C. conceptualized the study, performed experiments including data analysis and preparing figures, supervised the project, and wrote the manuscript.

---

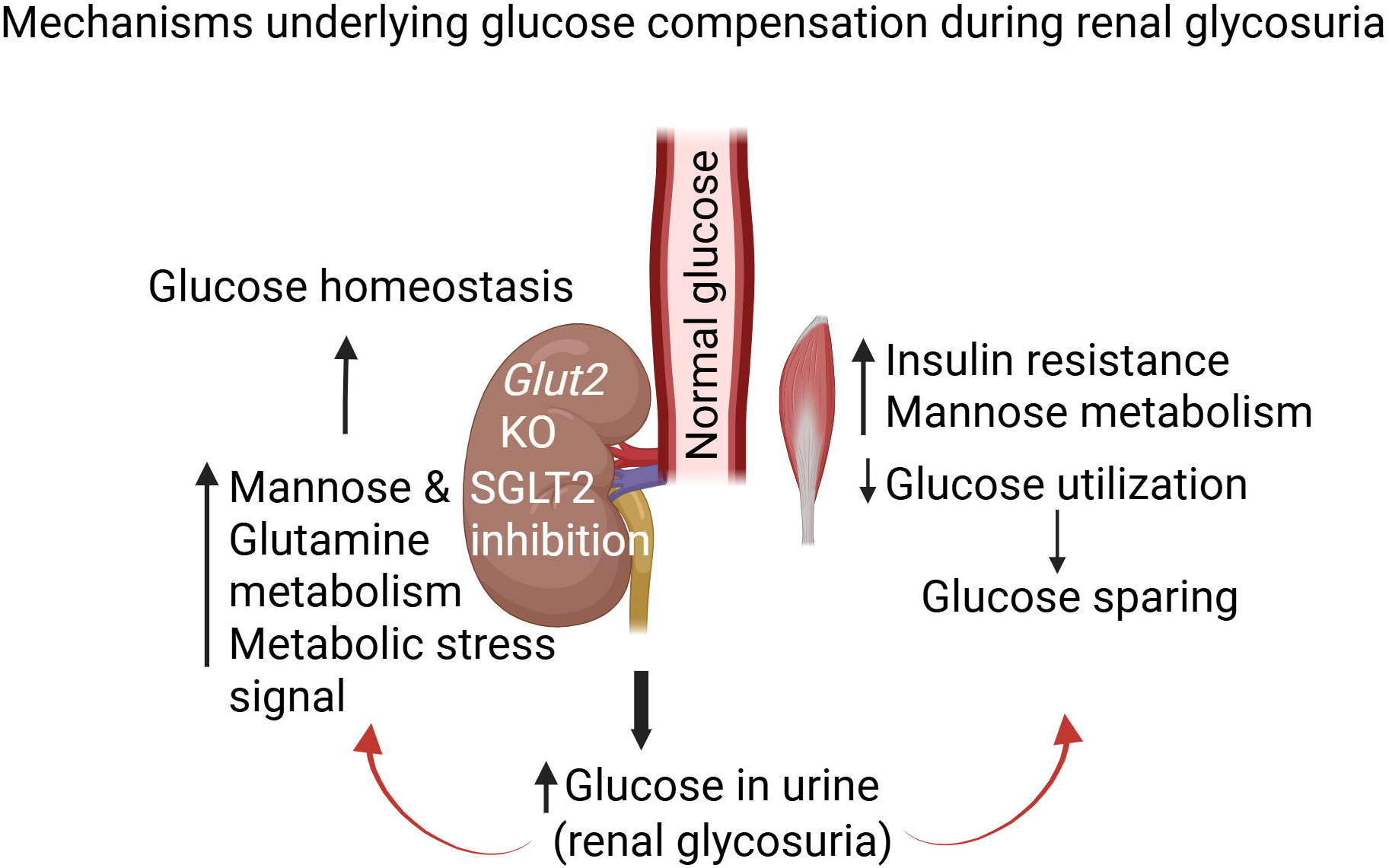

