## Supplementary figures for "Metabolic Reprogramming Coordinates Mannose and Glutamine Metabolism to Maintain Glucose Homeostasis During Glycosuria"

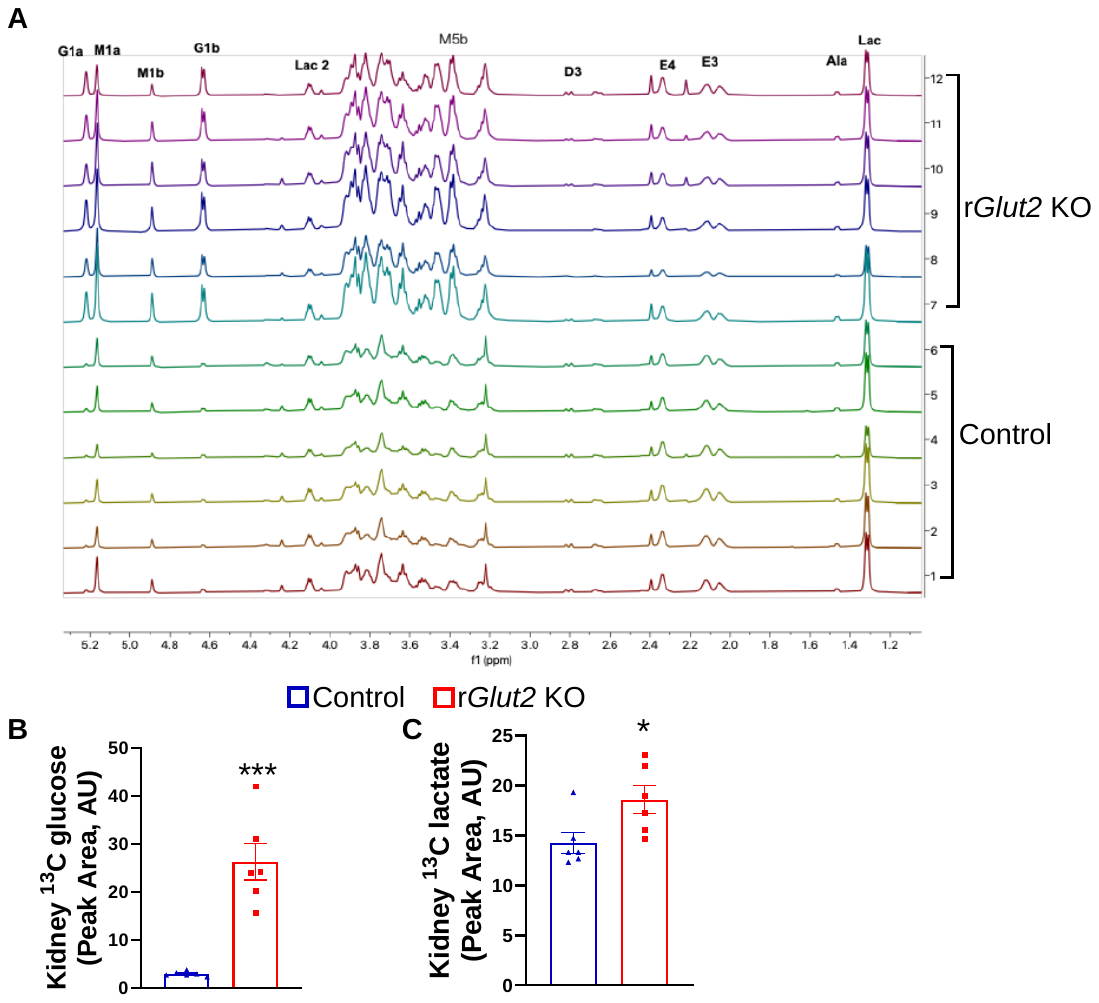

**Supplementary figure S1**: NMR analyses of kidney samples collected from 14-week-old male renal glucose transporter 2 knockout (r*Glut2* KO) mice and their control littermates 60 minutes after ^13^C_6_ mannose administration. **(A)** Representative NMR spectra; G, ^13^C glucose, M, ^13^C mannose, Lac, ^13^C lactate. The numbers following the metabolites represent the position of ^13^C atoms and the letter after that represents alpha and beta spin states of a nucleus. The X-axis represents parts per million. **(B)** ^13^C glucose and **(C)** ^13^C lactate in the kidney. Peak area normalized to d6-2,2-dimethyl-2-silapentane-5-sulfonate, AU, arbitrary unit. *p<0.05, ***p<0.001, two-tailed unpaired t-test. N=6.

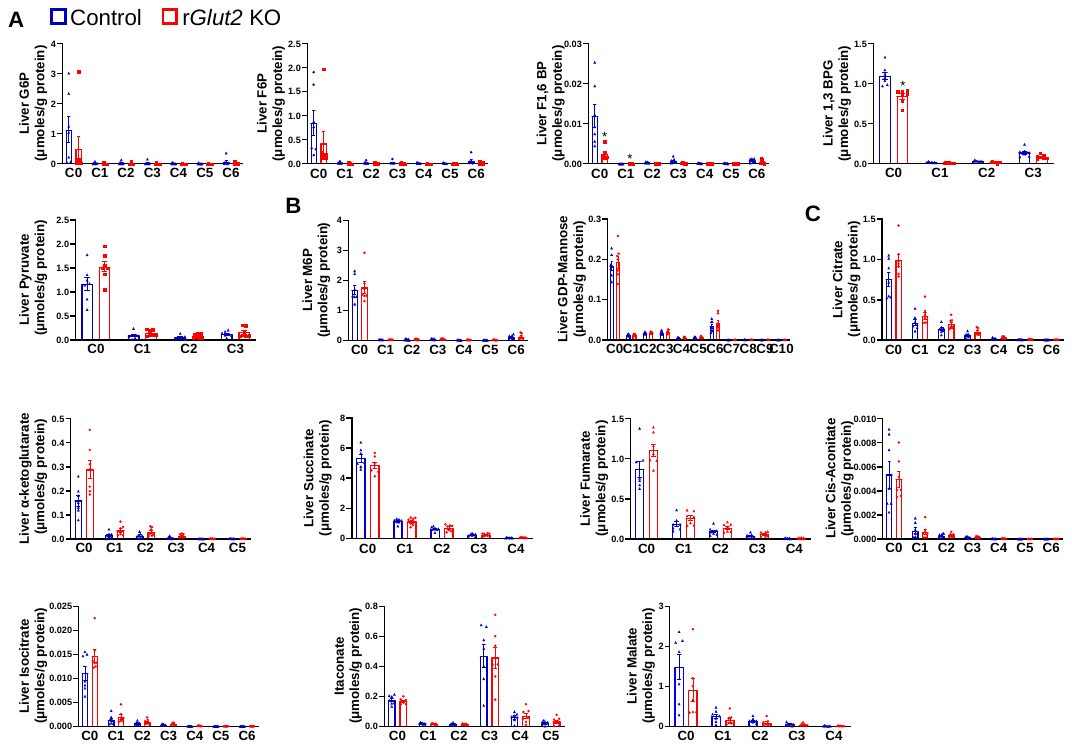

**Supplementary figure S2.** Stable isotope-resolved metabolomics analysis (by Ion Chromatography-Ultra-High Resolution Fourier Transform Mass Spectrometry) of liver glucose and mannose metabolism in 14-week-old male renal glucose transporter 2 knockout (r*Glut2* KO) mice and their control littermates 60 minutes after ^13^C_6_ glucose administration. **(A)** The contribution of the ^13^C to metabolites such as glucose 6-phosphate (G6P), fructose 6-phosphate (F6P), fructose 1,6, bisphosphate (F1,6BP), and pyruvate, **(B)** mannose 6-phosphate (M6P) and GDP-mannose, **(C)** the TCA cycle metabolites in the liver is similar in both the groups of mice. The X-axis represents the isotopologue distribution (number of ^13^C atoms) of each metabolite. N=7.

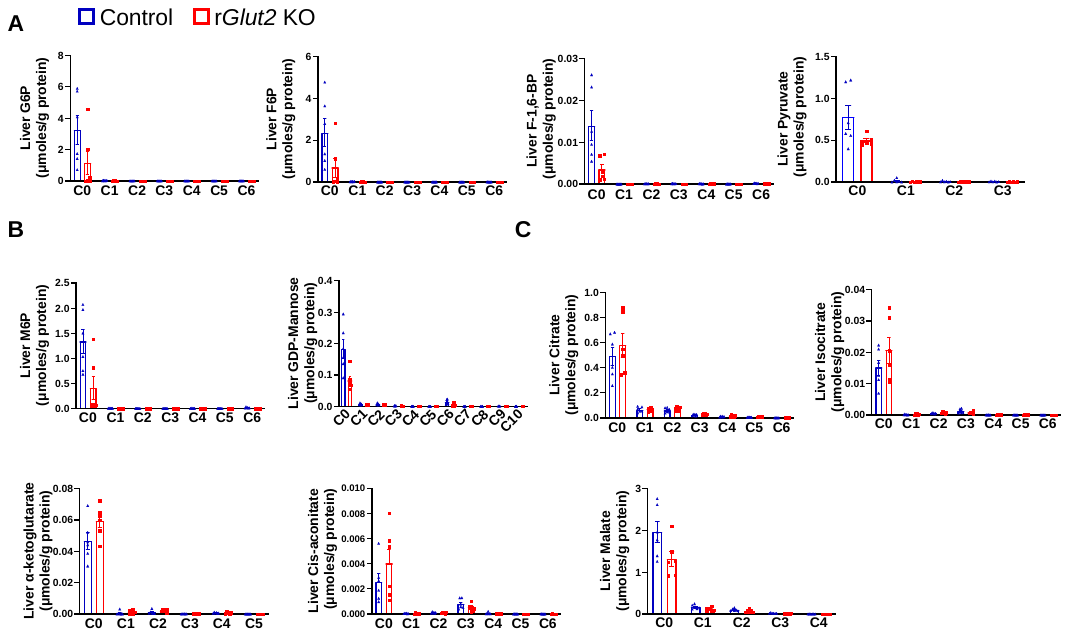

**Supplementary figure S3.** Stable isotope-resolved metabolomics analysis (by Ion Chromatography-Ultra-High Resolution Fourier Transform Mass Spectrometry) of liver glucose and mannose metabolism in 14-weeks-old male renal glucose transporter 2 knockout (r*Glut2* KO) mice and their control littermates 60 minutes after ^13^C_6_ mannose administration. **(A)** The contribution of the ^13^C to metabolites such as glucose 6-phosphate (G6P), fructose 6-phosphate (F6P), fructose 1,6, bisphosphate (F1,6BP), and pyruvate, **(B)** mannose 6-phosphate (M6P) and GDP-mannose, **(C)** the TCA cycle metabolites in the liver is similar in both the groups of mice. The X-axis represents the isotopologue distribution (number of ^13^C atoms) of each metabolite. N=7.

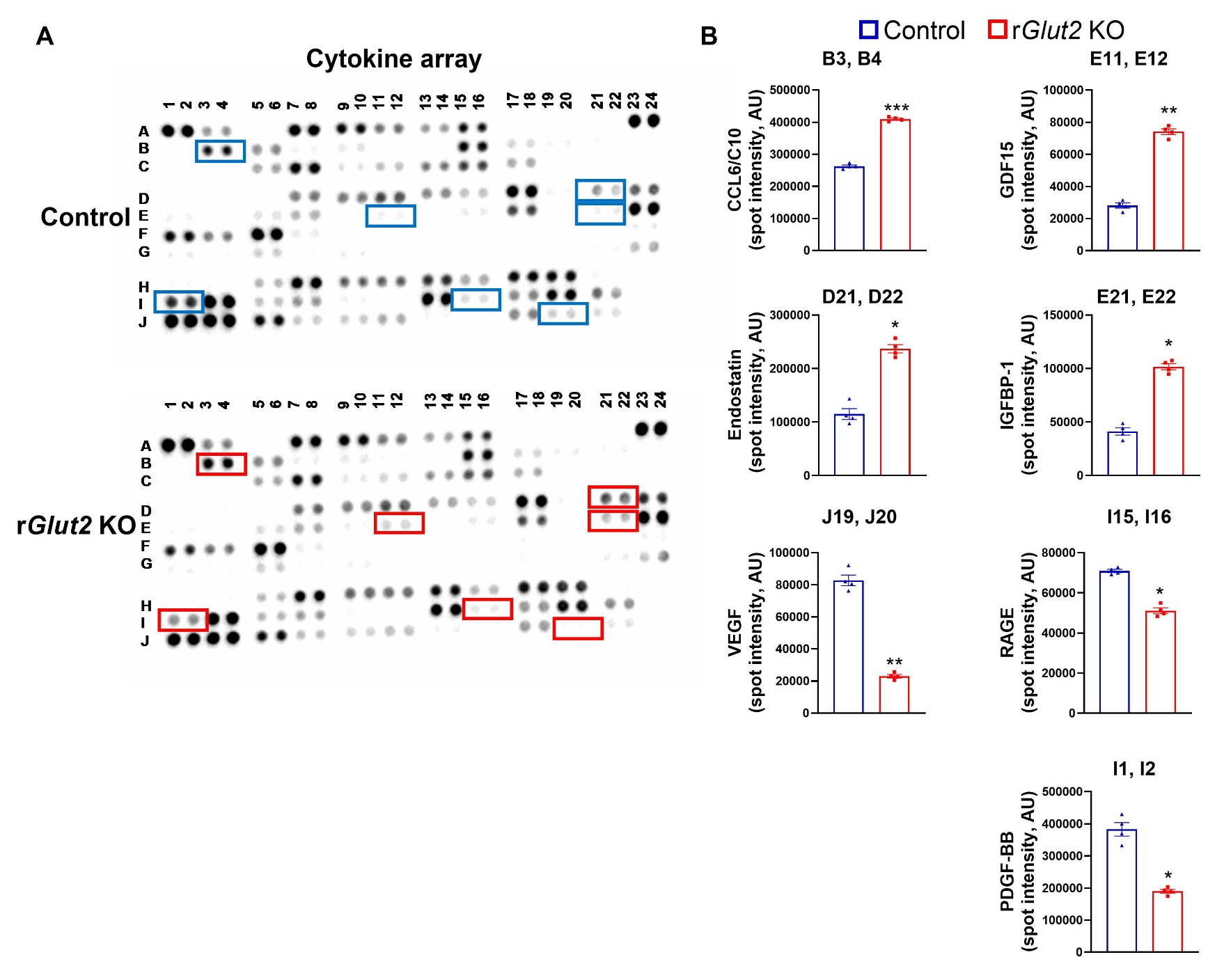

**Supplemental figure S4.** Cytokine array analysis of serum of 12-week-old male renal glucose transporter 2 knockout (r*Glut2* KO) mice and their control littermates. **(A)** Representative dot blot of the serum. **(B)** Graphs showing semi-quantification of the intensity of significantly changed spots (cytokines). The coordinates mentioned on each graph title represents the position of the corresponding immobilized antibody for a given cytokine on the array shown in **(A).** The increased cytokines represent metabolic‑stress signals elevated during renal glycosuria. N=4, 50 µl serum was pooled from three mice in each group, which was considered one sample; four such samples were prepared from 12 mice per group, *p<0.05, **p<0.01, two-tailed unpaired t-test. CCL6/C10, chemokine (C-C motif) ligand 6; GDF 15, growth differentiation factor 15; IGFBP-1, insulin-like growth factor binding protein 1; VEGF, vascular endothelial growth factor; RAGE, receptor for advanced glycation end-products; PDGF-BB, platelet-derived growth factor-BB.
